# Targeting the tomato fruit cuticle by gene overexpression and editing with the fruit epidermis-preferential *nsLTP* promoter

**DOI:** 10.1101/2024.11.11.622963

**Authors:** Marie Alonso, Ella Paulsen, Nathalie Geneix, Sandra Pedemay, Angelina D’Orlando, Didier Marion, Marc Lahaye, Bénédicte Bakan, Christophe Rothan, Johann Petit

## Abstract

The thick cuticle covering and embedding epidermal cells of tomato (*Solanum lycopersicum*) fruit, a model for cuticle studies, plays important roles in fruit protection and quality. To further our understanding of the influence of cuticle components on cuticle architecture and properties, the next step is to engineer cuticle composition. Ideally, to avoid indirect effects at whole-plant level, a fruit-and epidermis-specific promoter should be used. Here, to overexpress and edit target genes in tomato, we selected a *non-specific Lipid Transfer Protein 2* promoter (*pronsLTP*) with preferential activity in fruit epidermis, as shown by NLS-GFP fluorescence analysis. Overexpression of *SlMYB75* driven by *pronsLTP* induced anthocyanin accumulation specifically in the growing fruit epidermis, consistent with the cuticle deposition pattern. In keeping with changes in anthocyanins, the cuticle composition in flavonoids and phenolic acids was altered, as shown by Raman microspectroscopy, as were several cuticle properties. We then edited the carotenoid *PSY1* gene with a *pronsLTP*–driven *CRISPR/*Cas9 system. Different *PSY1* knock-out alleles were detected predominantly in fruit exocarp and mesocarp of color-impaired *pronsLTP::Cas9-Psy1* mutants, and very little in the leaf. A mutant carrying a heritable knock-out mutation in both fruit and leaf was also detected, indicating that mutants obtained with *pronsLTP* may require screening before being studied. Altogether, our results indicate that *pronsLTP2* can efficiently result in gene overexpression and editing in fruit epidermis. Implications of these findings are important for the functional analysis and genetic engineering of cuticle-related genes in tomato.

## Introduction

One of the most important goals of fruit growers is the production of high-quality fruit that can withstand transportation and storage. The fruit surface, which constitutes the interface between the fruit and its environment, acts as a protective barrier against pests, pathogens and water-loss. It therefore plays a crucial role in fruit shelf life and affects fruit appearance (color, glossiness, regularity, cracking…) and nutritional value^1^. In tomato, the fruit exocarp (also called peel or skin) is made up of an epidermal cell layer covered with the cuticle, and of several layers of sub-epidermal cells whose function is to generate mesocarp cells in the growing fruit^2,3,4,5^. The mesocarp underneath is a fleshy and heterogeneous tissue consisting of large polyploid cells which accumulate water, soluble sugars and organic acids^2,5^. The last cell layer of the pericarp facing the locular tissue is the endocarp. It consists of a single inner epidermal cell layer topped by the inner cuticle, both of which share common features with the outer epidermis and outer cuticle^6^. In addition to being a fleshy fruit model for which a large collection of genetic and genomic resources and tools is available^7^, tomato recently emerged as a model for cuticle studies. Its thick and astomatous cuticle is easy to peel and can be studied using a wide range of chemical and physical technologies^1,8,9,10,11,12^. In addition, the availability of tomato natural and artificially-induced cuticle mutants and transgenic lines^13^ has enabled the discovery of new cuticle-related functions and mechanisms^9,14,15^.

The fruit cuticle is covered by an epicuticular film of waxes which are also embedded within the cutin layer. Waxes are a complex mixture of derivatives of very-long-chain fatty acids (VLCFA), mainly alkanes and alcohols, and also includes various secondary metabolites, such as amyrins, flavonoids and sterols^16^. The main component of the cuticle is cutin^17^, which forms a continuum with cell wall (CW) polysaccharides of epidermal cells^18,19^. Cutin is mainly constituted by a polyester of polyhydroxy and epoxy fatty acids and fatty alcohols, and by varying amounts of phenolic compounds (e.g. p-coumaric acid and flavonoids)^12^. Like waxes, with which it shares common precursors, cutin precursors synthesized in epidermal cells are exported to CW through ABC transporters^15^. In the CW, cutin precursors are assembled by cutin synthases into a network of linear and branched cutin polymers^9,17,19^. A growing body of evidence suggests a role for nsLTP proteins in the transfer and deposition of the lipid monomers required for cutin assembly in the cell wall^20^.

Fruit skin properties largely depend on the composition and structure of the cuticle^8,12^. Lipid polyesters interact with non-lipid components including cuticle-embedded cell wall polysaccharides (CEP) and phenolics to determine cuticle architecture and properties^12,21,23^. CEP are diverse (cellulose, hemicelluloses, pectins) and exhibit specific features such as a high degree of pectin esterification (methylation and acetylation) and a high cellulose crystallinity^14,18^. Very recently, it has been shown that the growing fruit cuticle undergoes compositional and macromolecular rearrangements such as the specific spatiotemporal accumulation of phenolic compounds (p-coumaric acid and flavonoids) and the dynamic remodeling of CEP^21,23^. Furthermore, advanced technologies including confocal Raman microspectroscopy and Peak-Force Quantitative Nanomapping (PF-QNM) mode of AFM allowed us to show that distinct mechanical areas are correlated with the chemical and structural gradients in the cuticle^23^.

Altogether, these findings suggest that CEP and phenolics fulfill distinct functions in the cuticle during fruit development. The question now is to decipher how these compounds can modulate cuticle architecture and properties. Tomato cuticle mutants^13^, including *cutin-deficient* mutants^4,11,14,19,24,25,26,27^, have been instrumental in advancing our understanding of cuticle formation. Likewise, overexpression, RNAi silencing or CRISPR/Cas9 gene editing of CEP and phenolics-related genes in epidermal cells should bring insights into interactions between cutin, polysaccharides and phenolics. However, in contrast to cuticular lipid genes which are specifically expressed in epidermal cells^6,28^, many CW and phenolic-related genes are also expressed in various fruit tissues and plant organs^28^ where they fulfil important roles. Therefore, to avoid undesirable effects at plant and fruit levels, and their possible repercussions on the fruit cuticle, genes-of-interest should be targeted specifically to the fruit exocarp at times corresponding to the formation of the cuticle.

Here, we selected the promoter of a cuticle-related *non-specific Lipid Transfer Protein* (*nsLTP*) gene highly expressed in the fruit epidermis for its potential use for cuticle studies in tomato fruit. To this end, we assessed its suitability to drive specific gene expression in exocarp cells of early developing tomato fruit. Findings show that stable ectopic expression of the *Sl*MYB75 transcription factor under the control of *pronsLTP* leads to preferential alterations of the phenylpropanoid pathway and/or anthocyanin accumulation in epidermal cells of the fruit, thereby changing cuticle composition, assessed by Raman microspectroscopy, and properties. These results indicate that *pronsLTP* is well suited for overexpressing target genes preferentially in the fruit epidermis, thus enabling the study of the contribution of lipids, polysaccharides and phenolics to cuticle architecture and properties. Moreover, CRISPR/Cas9 gene edited mutants of *PHYTOENE SYNTHASE 1* (*PSY1)*, chosen as a visual marker of fruit color, displayed high mutation frequency in the fruit exocarp and mesocarp, and very little in the leaf, indicating that *pronsLTP* is suitable for efficient mutagenesis of fruit cuticle-related target genes in tomato.

## Results and Discussion

### The cuticle-related *non-specific Lipid Transfer protein 2* (*nsLTP2)* gene is preferential to the fruit exocarp

Cuticular material is mainly deposited into the outer epidermal cell wall at early stages of fruit development^21,24^ while sub-epidermal cells may also synthesize smaller amounts of cuticular components^21,27^. A promoter suitable for engineering the fruit cuticle without affecting other fruit tissues or plant organs should therefore meet certain requirements. Ideally, such promoter should (i) be specific to the exocarp and/or epidermis of the fruit; (ii) follow a specific temporal expression during fruit development, in line with cuticle formation that is maximum during fruit growth; (iii) and, of course, be strong enough to efficiently drive transgene expression in the fruit epidermis. Following these guidelines, we excluded the *proCRC*, *proTPRP* and *proPG* fruit-specific promoters we had previously analyzed in the miniature cultivar Microtom^29^, as well as the ripening-specific *E8* promoter^30^. The most efficient fruit promoter targeting early stages of fruit development in tomato is the widely used *proPPC2*^29,31,32^. Unfortunately, *proPPC2*, which drives a strong expression in mesocarp cells, is not active in epidermal cells^29^. Promoters from the tomato *SlCER6* gene encoding a VLCFA elongase β-ketoacyl-CoA synthase involved in wax biosynthesis and *SlCHS1* gene encoding a chalcone synthase enzyme involved in flavonoid biosynthesis were shown to drive reporter gene expression in fruit exocarp^3^. However, the *SlCER6* gene is expressed in both leaves and fruit and its mutation has considerable effects on leaf wax load and composition^33^ while the *SlCHS1* gene is also strongly expressed in flower buds according to the Tomato eFP browser Rose Lab Atlas at (http://bar.utoronto.ca). Very recently, the promoter of the tomato *PATHOGENESIS RELATED PROTEIN 10* (*SlPR10*) gene has been shown to drive a strong and preferential expression of the SlANT1 MYB transcription factor (TF) in the fruit exocarp, which resulted in strong anthocyanin accumulation in the fruit skin^34^. Moreover, fruit expressing the wax-regulating MYB31 TF under the control of *proPR10* exhibited alterations of cuticle properties, thus making *SlPR10* promoter very attractive for fruit cuticle engineering. However, analysis of transgenic plants where *SlANT1* expression was driven by the *SlPR10* promoter showed that it is also active in other plant organs, including leaf and stem^34^, which may impede its use when gene overexpression or silencing are lethal or have detrimental effects on the plant.

To identify additional candidate promoters, we used tomato gene expression data previously reported for fruit skin/exocarp^2,3^ or specific fruit tissues or cell-types^6,28^. Mining the semi-quantitative gene expression data obtained by Laser-Microdissection/RNAseq analysis (LMD-RNA seq) of fruit tissues and cell-types at the fruit expansion stage, showed that 15 out of the 25 top hits for genes expressed in the outer epidermis are *nsLTP* genes (Table S1). These include the strongly expressed *nsLTP1* and *nsLTP2* genes encoding well known allergens found in tomato peel^35^. As the nsLTP proteins may be involved in cuticle formation but also fulfil other functions in the plant, for example in plant defense^28,36^, we examined with the Tomato eFP browser whether the *nsLTP1* and *nsLTP2* genes were specifically expressed in the fruit. To determine if *nsLTP1* and *nsLTP2* are preferentially expressed in the epidermal and sub-epidermal cell layers at early stages of fruit development, we further searched the tomato fruit gene atlas (TEA-SGN database at https://tea.solgenomics.net) obtained by quantitative LMD-RNA seq analysis of fruit tissues and cell-types throughout fruit development^28^. TEA-SGN can therefore be mined for spatiotemporal expression patterns of candidate genes and clusters of co-expressed genes in the fruit.

Remarkably, most genes co-expressed with *nsLTP1* and *nsLTP2* genes (r>0.7), i.e. genes displaying similar expression patterns in the various pericarp cell-types along fruit development, could be ranged in two categories (Table S2). The first category includes genes involved in cuticle-related pathways including wax (*LACS*, *KCS6*, *CER26*, *CER1*) and cutin (*LACS*, *CYP77A2*) biosynthesis; cutin export and assembly (*nsLTP, GELP, BODYGUARD)*; and their regulation (*MIXTA-like* transcription factor) (Figure S1). The second category includes genes involved in flavonoid biosynthesis and, more specifically, anthocyanin biosynthesis (*4CL*, *CHS*, *CHI*, *F3H*, *DFR*, *ANS*, *F-3GT*) and its regulation (*MYB12* transcription factor) (Figure S2). In addition, there are genes likely to be involved in plant defense including *nsLTP* and *PATHOGENIS RELATED* protein 10 (*PR10*). Examination of gene expression in the whole plant using the Tomato eFP browser showed that all cuticle-related genes found in the cluster were not fruit-specific, with the exception of *nsLTP2*. Most flavonoid-related genes were found to be preferentially expressed in the fruit exocarp (Table S2), in agreement with^3^ but also, for the most highly expressed ones e.g. *CHS2*, in the flower bud.

We therefore focused our study on the *nsLTP2* gene (*Solyc10g075090*), which has high and preferential expression in both inner and outer fruit epidermis, from 5 DPA to the Mature Green stage, i.e. when cuticle deposition rate is the highest^3,24^. In addition, unlike *CHS2* and *PR10*, *nsLTP2* transcript abundance is, by far, much lower in the collenchyma (sub-epidermal cells) and parenchyma (mesocarp cells) (Table S2) allowing preferential targeting of inner and outer cuticles. To further explore the link of *nsLTP2* with cutin deposition, we next analyzed its expression in *cutin-deficient* and *cutin-enriched* tomato mutants^24,25^. In the *cus1* RNAi-silenced Wva106 tomato (cherry) transgenic lines obtained previously^9^, we observed a remarkable correlation between the abundance of *CUS1* and *nsLTP2* transcripts (Figure 1A). We found the same relationships between the abundance of *nsLTP* and *CUS1* transcripts in the Micro-Tom *cutin-deficient Slcus1* and *Slgpat6* and *cutin-enriched* P23F12 mutants previously described^13,24,25^ (Figure S1), where *nsLTP2* transcript abundance was correlated with that of *CUS1* (Figure 1B). These results, which suggest a complex yet unexplored relationship between *nsLTP2*, cutin deposition and *CUS1* expression, strengthen the potential suitability of the *nsLTP2* promoter to drive the overexpression and possible editing of target genes in the fruit epidermis. Such tool should enable the exploration of the contribution of cutin-embedded polysaccharides, phenolics and flavonoids in cuticle architecture and properties^12^.

**Figure 1.**
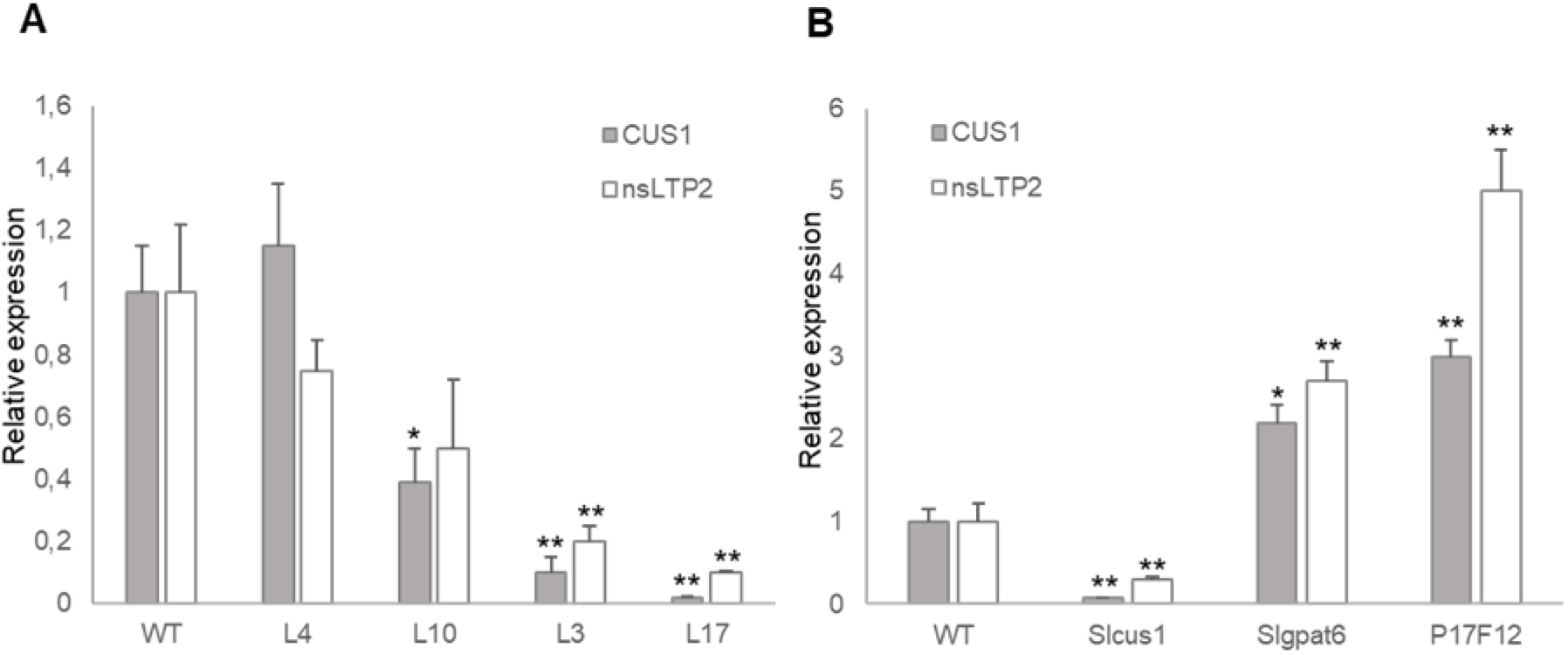
*nsLTP2* expression in *cuticle-deficient* and *cuticle-enriched* tomato mutants. Comparative RT-qPCR analysis of *SlnsLTP2* and *SlCUS1* transcript levels in (A) *cus1* RNAi transgenic lines L4, L10, L3 and L17; (B) *cuticle-deficient* EMS mutants *cus1* and *gpat6,* and *cuticle-enriched* EMS mutant P17F12. Values are mean ± SD (*n* = 3). *, *P* < 0.05; **, *P* < 0.01 (Student’s *t* test).

### The *nsLTP* promoter drives *GFP* expression preferentially in fruit exocarp

To date, various reporter genes have been used in tomato to study the spatio-temporal activity of fruit-specific promoters, including green fluorescent protein (GFP) and its fluorescent derivatives, luciferase-based luminescence and GUS^3,29,31,34,37^. Here, we used GFP tagged with the nuclear localization signal (NLS)^29^ as a reporter gene to study the activity of the *nsLTP2* promoter (*pronsLTP*) in vegetative organs and fruit (Figure 2). Observation of GFP fluorescence highlighted the bright green color of fruit surface until 20 DPA, which was less intense at the Mature Green (MG) stage, before the onset of ripening (Figure 2A). No fluorescence was detected in wild-type (WT). Examination of the 20 DPA fruit cross-section showed that the bright green color was clearly restricted to the fruit skin; a faint green color was also observed in the seeds (data not shown). Consistent with the distribution of *nsLTP2* transcripts in the plant (Tomato eFP browser), no *pronsLTP* activity was detected in other plant organs e.g. flower and root (Figure 2B), indicating that no side-effects on other plant organs should be expected when using *pronsLTP*. Unexpectedly, however, a bright green color was seen around the leaf veins (Figure 2B) and, more specifically, in the trichomes of various types that are enriched in this area (Figure 2B, Figure S3). This very localized GFP expression indicates that *pronsLTP* could also be used for the production of specialized metabolites in tomato trichomes^38,39^. To further analyze the spatial distribution of GFP in the diverse cell types of the exocarp of *pronsLTP*::NLS-GFP, we performed confocal analysis of 5 DPA fruit cross-sections stained with calcofluor to visualize cell walls^40^ (Figure 2C).

**Figure 2.**
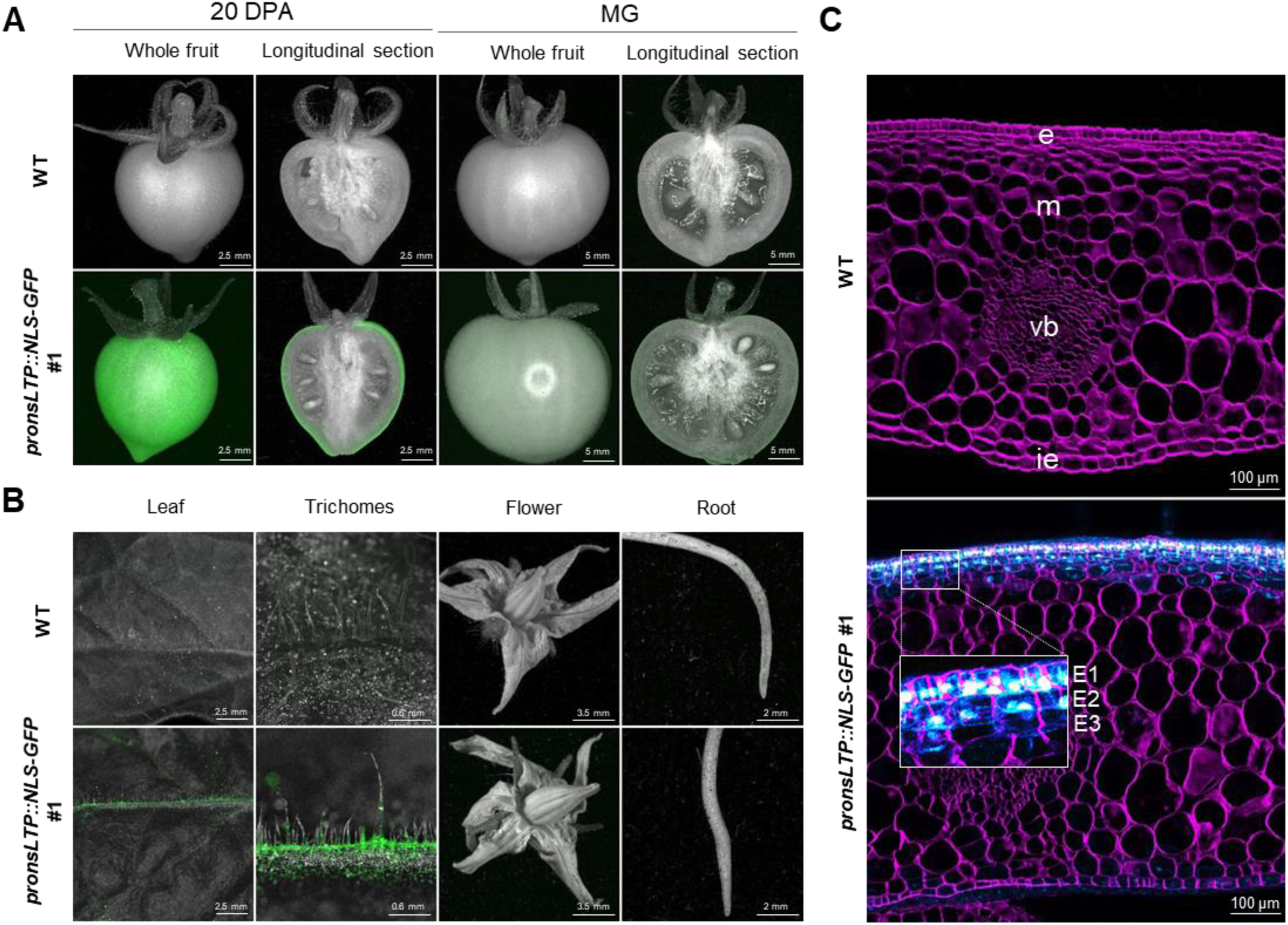
Detection of *pronsLTP*-driven nuclear GFP in tomato fruit and leaf trichomes. (A) Macroscopic detection of GFP fluorescence of *pronsLTP::NLS-GFP* fruit. Overlays of black-and-white and GFP images are shown for representative 20 DPA and MG fruits of *pronsLTP::NLS-GFP* #1 and WT. Pericarp GFP fluorescence is restricted to the exocarp. DPA, Days Post Anthesis; MG, Mature Green. (B) Macroscopic detection of GFP fluorescence in leaf, trichomes, flower and root of *pronsLTP::NLS-GFP* #1 and WT. Overlays of black-and-white and GFP images. Leaf GFP fluorescence is restricted to trichomes. (C) Confocal microscopy detection of GFP fluorescence (in blue) after calcofluor staining (in purple) of cross section of 5 DPA fruit pericarp of WT and *pronsLTP::NLS-GFP* #1. Specific GFP expression is seen in fruit exocarp and inner epidermis. Inset, magnified image of exocarp showing preferential localization of GFP in nucleus from outer epidermis (E1 cell layer), less intense fluorescence of nucleus in E2 sub-epidermal cell layer and few fluorescent nuclei in E3 cells. e, outer epidermis; m, mesocarp; vb, vascular bundle; ie, inner epidermis; E1, outer epidermis, layer 1; E2, sub epidermis, layer 2; E3, sub epidermis, layer 3.

Intense GFP fluorescence was observed in all cells of the outer epidermis (E1 cell layer) while the nuclei of sub-epidermal cells of the E2 layer also displayed GFP fluorescence as well as some sub-epidermal cells of the E3 layer (Figure 2C). A weak fluorescence was also seen in cells of the inner epidermis but no fluorescence was observed in the enlarging cells of the mesocarp, which is consistent with TEA-SGN data. These results indicate that *pronsLTP* is likely well suited to study genes contributing to cuticle formation in the developing fruit, in both the inner and outer epidermis of the fruit.

### Overexpression of *SlMYB75* under the control of *pronsLTP* alters the color of the fruit exocarp

Systems based on the visual aspect of the fruit are particularly attractive to as they do not require dedicated (fluorescence, luminescence) or destructive (GUS) technologies. Tomato fruit color has been successfully engineered by mutating genes involved in carotenoid biosynthesis (*PHYTOENE DESATURASE PDS* and *PHYTOENE SYNTHASE PSY*) and regulation of the flavonoid pathway (*MYB12*) ^41,42,43,44^; or by the ectopic expression of transcription factors (TFs) triggering the accumulation of anthocyanins such as Ros/Del TFs from snapdragon^45^ and tomato ANT1^34^ and MYB75^46^. Here, we chose the overexpression (OE) of tomato *SlMYB75* to provide visual, non-destructive, monitoring of *nsLTP* promoter activity in plant organs and fruit tissues. Furthermore, by enhancing the phenylpropanoid flux towards the synthesis of flavonoids, some of which are exported to the cuticle^3,8^, and anthocyanins, which are sequestered in the vacuole (Figure S1), we may expect changes in phenolic and flavonoid composition of the cuticle. This, in turn, could have an impact on cuticle architecture and properties^8,12^.

We therefore generated *pronsLTP::MYB75* and *pro35S::MYB75* lines (control), using established protocols for the transformation and selection of diploid transformants in the miniature MicroTom cultivar^29^. Between 12 and 15 independently transformed plants were generated for each *pro::MYB75* transgene construct. Following visual inspection of fruit color in T0 transformants, 2 representative lines were selected for each construct. The *SlMYB75*-induced purple color pattern of 6 kanamycin-resistant T1 plants per line was then characterized in the whole plant, leaf and open flower and throughout fruit development at 5 DPA, 10 DPA, 15 DPA, 20 DPA, mature green (MG; transition to ripening), and red ripe (RR; ripe fruit) stages (Figure 3A). As expected, a pronounced purple color was observed in all *pro35S::MYB75* plant organs, while no staining was detected in WT controls (Figure 3A). However, color distribution was more uneven than the uniform and intense purple skin color previously reported for *SlMYB75* overexpression in tomato^46^ and more similar to the activation-tagged *ANT1* line^47^. Likely explanations are the environmental conditions prevailing in our greenhouse, as the MYB-triggered anthocyanin biosynthesis is very sensitive to stress conditions and, particularly, to light stress^48^. The *pronsLTP* promoter activity was restricted to the fruit, as no purple color was observed in vegetative organs nor in flowers (Figure 3A). Consistent with the *nsLTP* transcript accumulation in the fruit (Table S2), *pronsLTP::MYB75* fruits displayed a weaker but consistent purple color at early stages of fruit development, from 5 DPA to 20 DPA, with a maximum from 5 DPA to 15 DPA. Additionally, a brown color was observed in *pronsLTP::MYB75* seeds at RR fruit stage, as in *pro35S::MYB75* seeds (Figure 3A), indicating the deposition in the seed coat of flavonoid polymers which, when oxidized, cause the seed to turn brown^49^. This activity is consistent with *nsLTP2* expression in the seed, which is restricted to the tegument at early stages of fruit development (Tomato Seed eFP Browser at http://bar.utoronto.ca). In the internal part of the *pro35S::MYB75* fruit, the purple color was observed in all tissues including pericarp, columella, placenta and locular tissue while only the outer part of the pericarp displayed a weak purple color in *pronsLTP::MYB75* fruit (Figure 3A).

**Figure 3.**
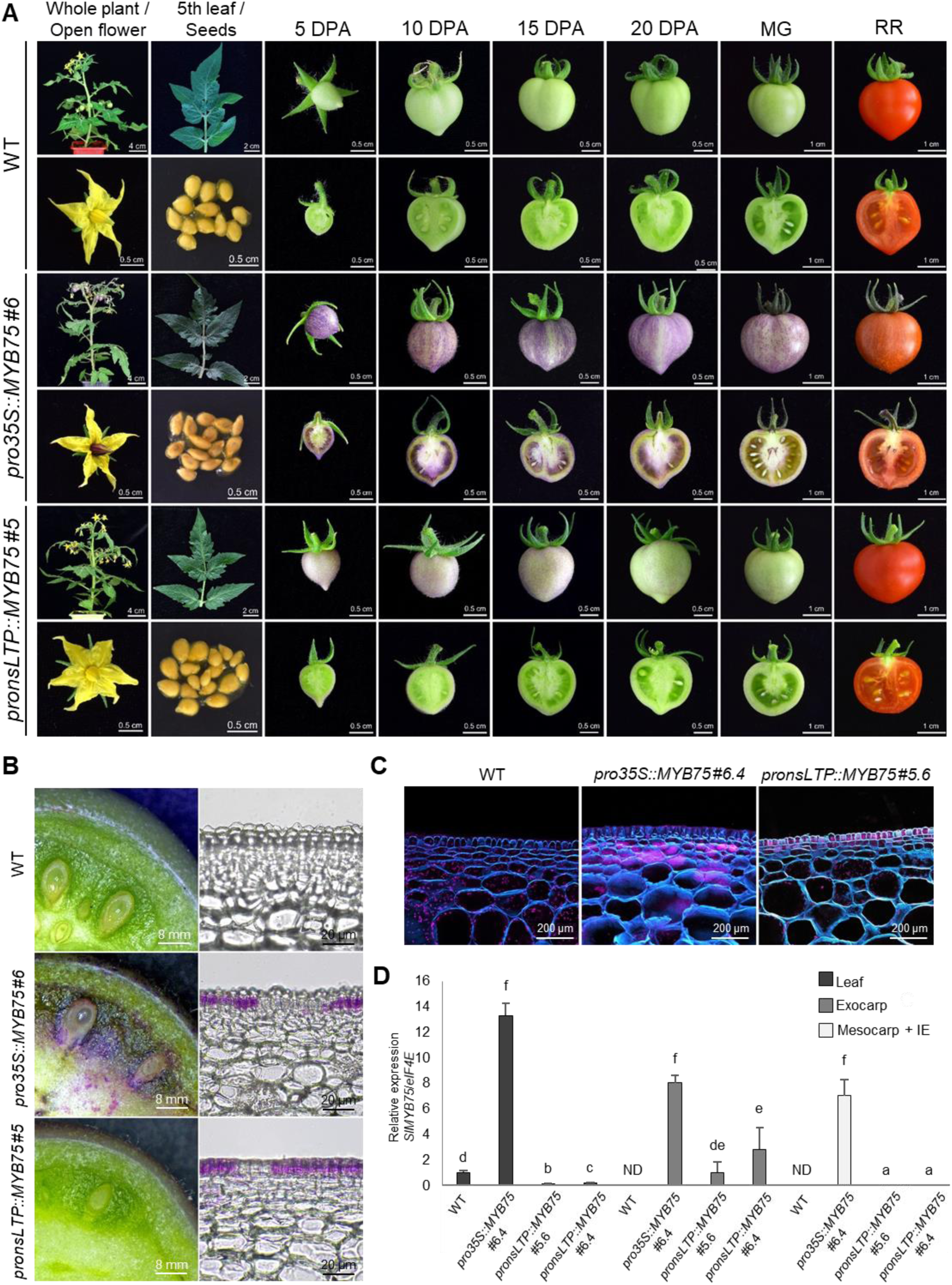
*pronsLTP*-driven expression of *SlMYB75* induces specific accumulation of anthocyanins in fruit exocarp. (A) Anthocyanin accumulation in WT, *pronsLTP::MYB75* and *pronsLTP::MYB75* transgenic lines. Whole plant, leaf, flower, seeds and developing fruit from 5 DPA to RR stage are shown. DPA, Days Post Anthesis; MG, Mature Green; RR, Red Ripe. (B) Distribution of anthocyanin accumulation in 10 DPA fruit of WT, *pro35S::MYB75* #6 and *pronsLTP::MYB75* #5 transgenic lines. Left: cross-section of the whole fruit; right: vibratome cross-section of the pericarp under the optical microscope. (C) Confocal microscopy detection of anthocyanins (purple) after calcofluor white staining (blue) of vibratome cross-section of 10 DPA fruit pericarp of WT, *prons35S::MYB75* #6 and *pronsLTP::35S* #5. Anthocyanins accumulate mostly in large mesocarp cells in *prons35S::MYB75* fruit, and in E1 and E2 exocarp cells in *pronsLTP::35S* fruit. Red dots in the cells are due to chloroplast autofluorescence. (D) RT-qPCR analysis of *SlMYB75* transcript abundance in leaf, fruit exocarp, and fruit mesocarp (including inner epidermis) of WT, *pro35S::MYB75* #6.4 and *pronsLTP::MYB75* #5.6 and #6.4. Values are mean ± SD (*n* = 3), with transcripts normalized to *eIF4E*. Lower-case letters indicate significant differences (P < 0.05; Kruskal-Wallis test with Tukey post-hoc test). ND, not detected.

Light microscopy examination of the distribution of purple color in cross-sections of 20 DPA fruits indicated that in *pronsLTP::MYB75* fruits, purple color was indeed restricted to the exocarp, unlike in *prons35S::MYB75* fruits (Figure 3B). Exocarps of both *pronsLTP::MYB75* and *prons35S::MYB75* fruit exocarp displayed an uneven distribution of purple color (Figure 3B), consistent with the variegated color of the fruit surface (Figure 3A). An intriguing feature was also the apparent lack of anthocyanin accumulation in the E1 cell layers of the *prons35S::MYB75* fruit (Figure 3B). Confocal microscopy observation of cross-sections of 20 DPA fruit from *prons35S::MYB75* stained with Calcofluor white (Figure 3C) further highlighted the large accumulation of anthocyanins in vacuoles of expanding mesocarp cells, as well as in E2 and E3 sub-epidermal cell layers but seemingly not in the E1 cell layer, in line with light microscopy observations (Figure 3B). In contrast, in *pronsLTP::MYB75* fruit, the anthocyanin accumulation was limited to the epidermis (E1) and E2 sub-epidermal cell layers. Moreover, epidermal cells of both *pro35S::MYB75* and *pronsLTP::MYB75* fruits displayed morphological alterations, as frequently seen in fruit cuticle mutants of various origins^4,8,26^. While E1 cells of WT exhibited a classical cone-shaped appearance, E1 cells from *pro35S::MYB75* were largely encased in thick CW/cuticle, which may explain their lack of anthocyanin accumulation. E1 cells of *pronsLTP::MYB75* were more square-shaped (Figure 3C).

Quantification by RT-qPCR of *SlMYB75* transcript abundance in plant and fruit tissues confirmed that the activity of the *nsLTP* promoter is limited to the fruit exocarp. In the *pronsLTP::MYB75* fruits, *SlMYB75* transcripts were detected only in the exocarp while in *pro35S::MYB75 fruits*, *SlMYB75* was found to be highly expressed in leaves and fruit mesocarp, in addition to fruit exocarp (Figure 3A).

### Raman microspectroscopy provides insights into the alterations in cuticle composition and properties

We further investigated if the *pronsLTP*-driven accumulation of anthocyanins in fruit exocarp could translate into modifications in cuticle composition, architecture or properties, as suggested by alterations in epidermal cell patterning (Figure 3C). We first analyzed water-loss and firmness of RR fruit and permeability to toluidine blue (TB) of MG fruit (Figure 4), all of which are associated with cuticle integrity. Such analyses could reveal cuticular defects as observed for example in the *cus1*, *gpat6* and *shn2* cutin-deficient mutants^24,25,26^. A slight ∼5.3 to 6.8% reduction in fruit fresh weight was detected in *pronsLTP::MYB75* fruits after 770 hours of storage at 30°C, depending on the line analyzed, but no differences in water-loss were seen in *pro35S::MYB75* fruits compared to WT (Figure 4A). While a trend towards reduced fruit firmness was observed for all *SlMYB75* overexpressing fruits analyzed, this reduction was significant only for *pronsLTP::MYB75#6.4* fruits (Figure 4B). The most remarkable visual alterations of cuticle properties were observed after fruit immersion in TB, which revealed large cracks in up to 50% of *pronsLTP::MYB75* fruits, the severity of which depended on the line (Figure 4C), and was consistent with the alterations in water loss and firmness observed (Figure 4A, 4B). Again, cuticle alterations were not observed in *pro35S::MYB75* fruits, indicating different cuticle behavior depending on the promoter used. This was unexpected given the greater accumulation of anthocyanins visible on the fruit surface in *pro35S::MYB75* fruits (Figure 2A).

**Figure 4:**
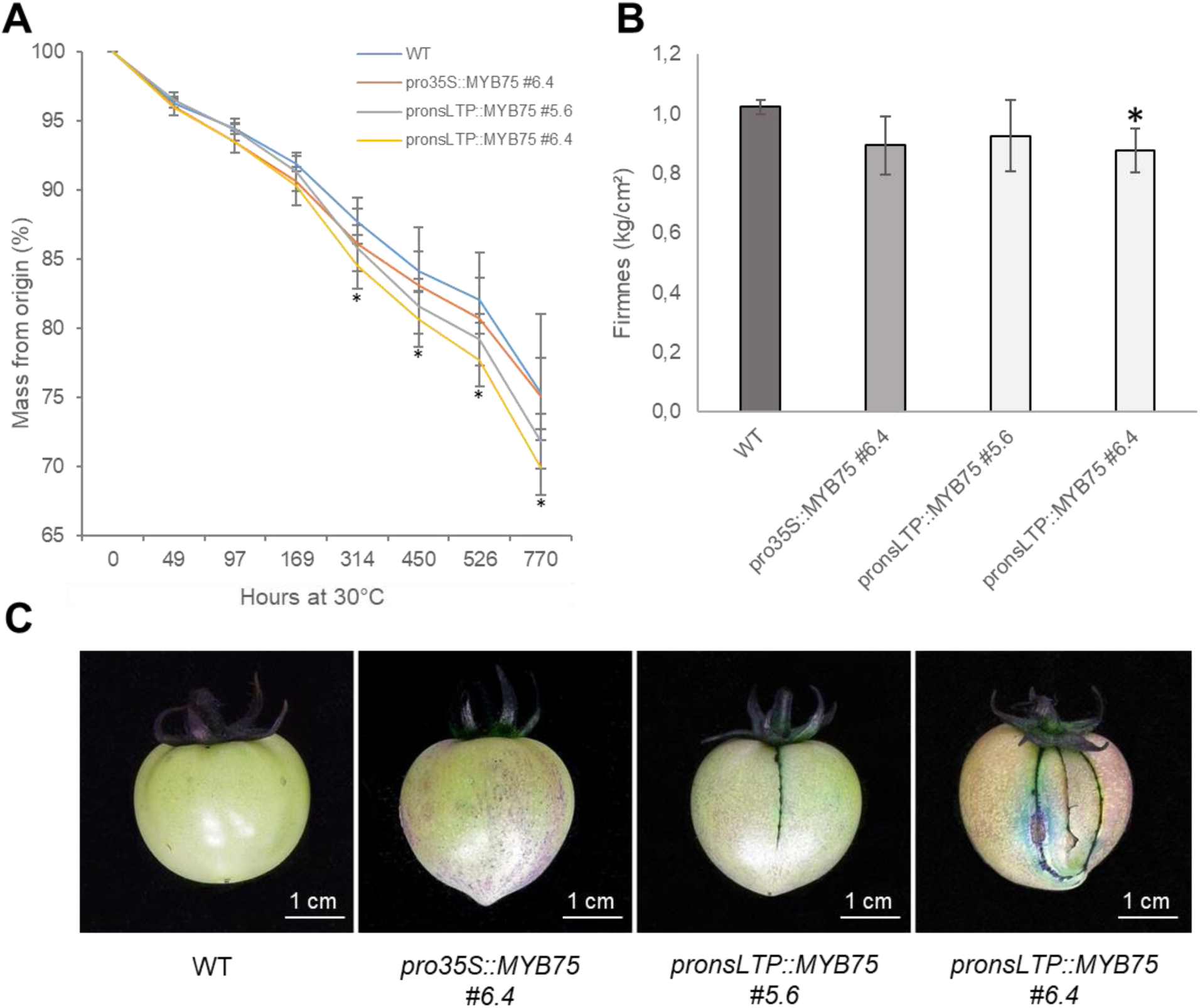
Cuticle properties of *SlMYB75* overexpressing fruits. (A) Water-loss in Red Ripe (RR) fruit of WT, *pro35S::MYB75 #6.4*, *pronsLTP::MYB75 #5.6* and *#6.4* during storage at 30°C. Masses were recorded at time zero (T0) and after 49, 97, 169, 314, 450, 526 et 770h. *, P < 0.05. (B) Penetrometry measurements on RR fruit. Values expressed in kg/cm² are mean ± SD (n = 5). *, P < 0.05; **, P < 0.01 (Student’s t test). (C) Permeability to toluidine blue of Mature Green fruits incubated with the dye during 16h. Photographs of representative fruits are shown.

To further investigate compositional changes possibly responsible for alterations in cuticle architecture and properties, we analyzed the fruit cuticle of transgenic lines overexpressing *SlMYB75* using an established Raman microspectroscopy technology that allows high resolution analysis of specific regions of the cuticle^21,22^. As we previously observed sharp chemical heterogeneities in the fruit cuticle^22^, we paid particular attention to always analyze in the same area i.e. at the junction of adjacent epidermal cells. To address the differences in fruit surface color observed in *pronsLTP::MYB75* and *pro35S::MYB75* fruits (Figure 3B), we analyzed both purple and white areas in MG fruit i.e. before the ripening-associated accumulation of naringenin and p-coumaric acid.

Raman spectral analyses combined with principal component analysis (PCA) clearly differentiated WT, *pronsLTP::MYB75* and *pro35S::MYB75* fruit cuticles. The Raman spectra acquired from purple zones of the cuticle (Figure 5A, 5B, 5C, 5D) of *pronsLTP::MYB75* and *pro35S::MYB75* fruits highlighted specific bands at 580 cm^-1^; 624 cm^-1^ and 1557 cm^-1^ assigned to flavonoids^21,50,51^, while the 1520 cm^-1^ band has been previously assigned to the *v*(C-C) band of anthocyanins with or without glucoside derivation^52,53^. This result, which indicates that the overexpression of *SlMYB75* triggered the accumulation of flavonoids, including anthocyanins, in the fruit epidermis, is in full agreement with the known role of *Sl*MYB75 in anthocyanin biosynthesis. Interestingly, *pronsLTP::MYB75* and *pro35S::MYB75* fruit cuticles seem also to contain less lipids according to the specific bands of CH2 (*τ*CH2 band at 1304 cm^-1^ and *δ*(CH2) band at 1440 cm^-1^) and less p coumaric acid (*v*(C=C) at 1605 cm^-1^)^21^, compared to the WT fruit cuticle. Raman spectra obtained from the white-colored zones of the fruits also demonstrated differences between the *SlMYB75* overexpressing lines and the WT, these alterations being more pronounced in *pro35S::MYB75* than in *pronsLTP::MYB75* (Figure 5E, 5F, 5G, 5H). Notably, the spectra of *pronsLTP::MYB75* and *pro35S::MYB75* fruits indicates a lower proportion of both lipids (1304 cm^-1^, 1440 cm^-1^) and phenolics (1167 cm^-1^, 1605 cm^-1^) including p-coumaric acid (Reynoud et al 2022). As the most abundant lipid found in the cuticle is the cutin polyester, these results suggest a reduction of cutin deposition in the *SlMYB75* overexpressing lines in addition to the changes in flavonoids and phenolics.

**Figure 5:**
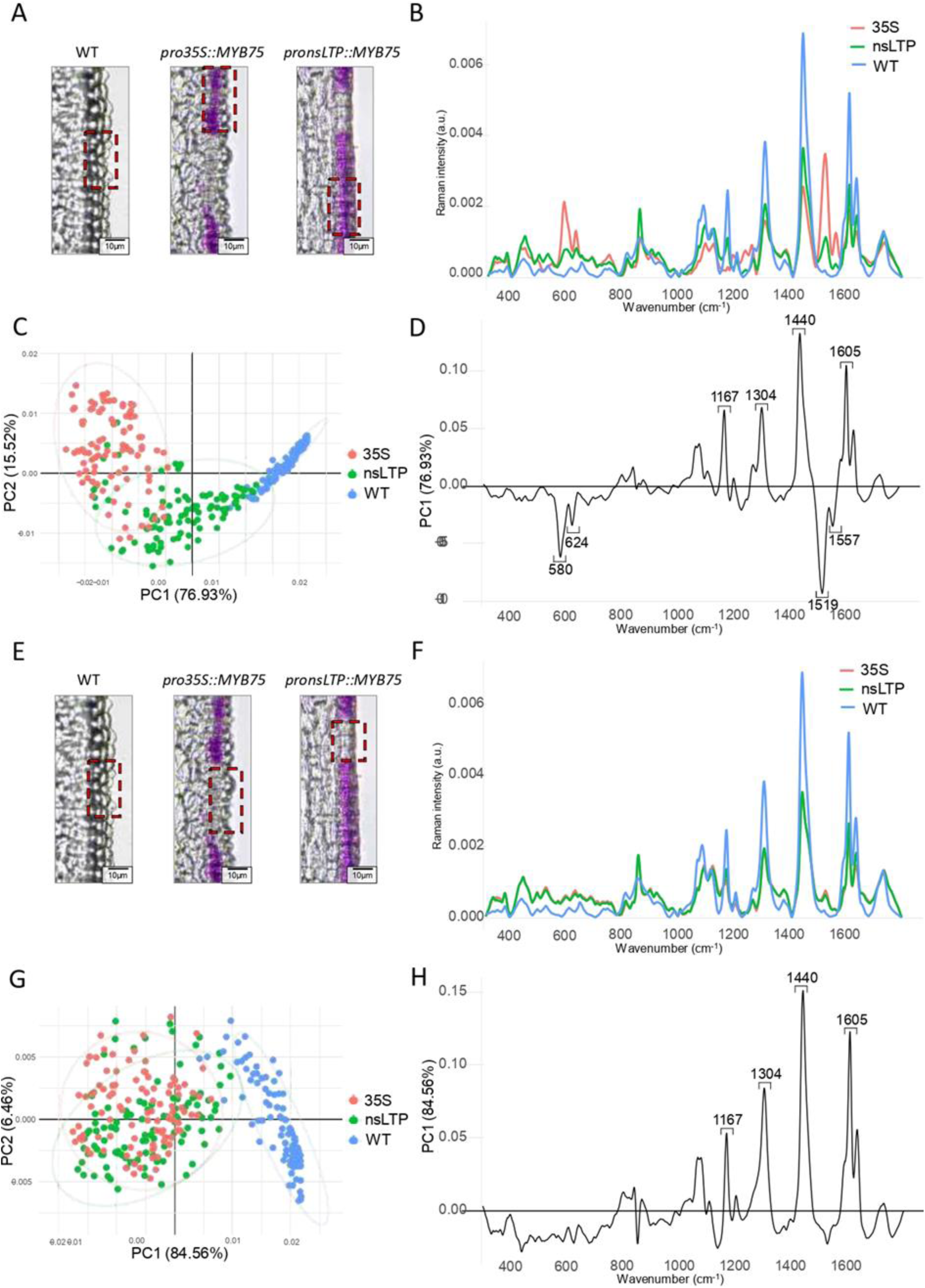
Raman spectroscopy analysis of the fruit cuticle composition of *SlMYB75* overexpressing fruits. (A) Areas of WT and purple epidermis analyzed by Raman spectroscopy in *SlMYB75* overexpressing lines. Photographs shown are from Figure 3B. (B) Average spectra acquired on WT and purple areas of *pro35S::MYB75 #6.4* and *pronsLTP::MYB75 #5.6.* Spectra were always taken between two adjacent epidermal cells. (C) Scatter plot of PCA analysis performed on spectra acquired from WT and purple areas of *pro35S::MYB75 #6.4* and *pronsLTP::MYB75 #5.6.* (D) PC1 loading of the PCA analysis shown in (C) discriminating the WT from the purple epidermis of *SlMYB75* overexpressing lines. (E) Areas of white epidermis analyzed by Raman spectroscopy in WT and *SlMYB75* overexpressing lines. Photographs shown are from Figure 3B. (F) Average spectra acquired on WT and white epidermis areas of *pro35S::MYB75 #6.4* and *pronsLTP::MYB75 #5.6*. Spectra were always taken between two adjacent epidermal cells. (G) Scatter plot of PCA analysis performed on spectra acquired from WT and white epidermis areas of *pro35S::MYB75 #6.4* and *pronsLTP::MYB75 #5.6*. (H) PC1 loading of the PCA analysis shown in (G) discriminating the WT from the white epidermis of *SlMYB75* overexpressing lines.

In the above analyses, anthocyanins were detected in the outer wall of epidermal cells, while these water-soluble compounds are sequestered in the cell vacuole. As anthocyanins may have been released during sample preparation and thus contaminated the cuticle, we sought to clarify specific changes in cuticle composition. To this end, cuticles were enzymatically isolated (to get rid of any non-cutinized material) and analyzed by RAMAN spectroscopy, including a step of normalization on the amount of lipids (Figure S4). The data obtained show a decrease in the relative content of phenolic compounds, in particular in p-coumaric acid, in *SlMYB75* overexpressing lines; these changes were also more pronounced in *pro35S::MYB75* than in *pronsLTP::MYB75*. Indeed, a 33 % reduction in the proportion of p-coumaric acid was observed in *pro35S::MYB75* while a 10% relative reduction in p-coumaric acid was observed in the *pronsLTP::MYB75* cuticle.

Altogether, our data indicate that both *pronsLTP::MYB75* and *pro35S::MYB75* induced specific deposition of anthocyanins-related compounds in the fruit epidermis, which was expected given the known role of *SlMYB75* in the regulation of the anthocyanin biosynthesis pathway^46^ and the expression of most genes belonging to the anthocyanin pathway in the fruit exocarp (Figure S2). Raman analyses of the fruit cuticle further showed that *SlMYB75* overexpression induced a reduction in cutin and the cutin-associated phenolic acid, p-coumaric acid, in the fruit cuticle. As p-coumaric acid is a precursor of flavonoids, including anthocyanins, this result likely indicates that the phenypropanoid flux is partly boosted towards the formation of anthocyanins, at the expense of the p-coumaric acid accumulation in the fruit cuticle. As *35S* and *nsLTP* promoters are both active in the fruit epidermis, according to the compositional changes observed, the difference in exocarp fruit color (purple vs white sectors) is intriguing. It possibly indicates that a required step in anthocyanin synthesis/glycosylation/storage is not being carried out in the white sectors, resulting in the formation of colorless anthocyanin precursors instead of colored anthocyanins.

Even more intriguing is the discrepancy between the visual appearance of the fruit (Figure 3) and the changes in cuticle composition induced by the *35S* and *nsLTP* promoters (Figure 5), in comparison with the severity of the alterations in cuticle properties (Figure 4). The mechanical properties of the cuticle, e.g. stiffness and extensibility, vary according to cuticle composition, developmental stage of the fruit and environmental conditions^54,55^. In particular, the cuticle should be able to deform to allow cell enlargement during the cell expansion phase, during which there is a considerable increase in fruit size^2^.

Increased anthocyanin biosynthesis induced by the ectopic expression of the Del/Ros TFs in tomato fruit was shown to affect wax accumulation^45^ but had no effect on the biomechanical properties of the tomato fruit cuticle^8,56^. Both naringenin chalcone, the predominant flavonoid, and p-coumaric acid, the major phenolic acid, accumulate during fruit ripening in the cuticle^8,21,22,56^, and may contribute to its stiffening^8^. Indeed, transient CHS silencing indicate that flavonoids stiffen the elastic phase and reduce permanent viscoelastic deformation of the cuticle^8,56^. Thus, the modifications in cuticle composition observed by Raman microspectroscopy of *SlMYB75* overexpressing fruit cuticles, which showed changes in flavonoids and p-coumaric acid content, are consistent with cracking, the main alteration in cuticle properties of *pronsLTP::MYB75* fruit. One possible explanation for the discrepancy observed when using the two promoters is that of the spatiotemporal expression pattern of *nsLTP*, which drives more specific and intense expression of *NLS-GFP* and *SlMYB75* in epidermal cells at early stages of fruit development, where the cuticle is mainly formed^24^, than the *35S* promoter (Figures 2, 3). These results highlight the interest of using the *pro35S::MYB75* and *pronsLTP::MYB75* lines generated here for deciphering the relationships between cuticle properties, architecture and composition e.g. by correlative multimodal imaging approach to analyze the relationships between cuticle composition and mechanical properties, as previously done^22^.

### The *nsLTP* promoter drives efficient gene editing of the carotenoid *PSY1* gene in the fruit exocarp

The red color of ripe tomato fruit is largely due to lycopene, a carotenoid compound which accumulates in both fruit flesh and peel. A key step in the ripening-associated carotenoid biosynthesis is catalyzed by the *PHYTOENE SYNTHASE 1* (*PSY1*) gene, whose expression is strongly induced at the onset of fruit ripening. Mutation in the N-terminal region of the PSY1 protein resulting in a truncated protein leads to a block of the carotenoid biosynthesis pathway and hence to yellow-orange colored tomato fruits^41^, whose residual color is due to the accumulation of the yellow-colored naringenin chalcone. We therefore targeted *PSY1* by the CRISPR/Cas9 system to provide a visual marker for efficient gene editing. As expected, gene editing of *PSY1* under the control of the 2X35S promoter (*pro35S::Cas9-Psy1*) produced yellow-colored fruits (Figure 6A), due to the strong reduction in the carotenoid content of the fruit pericarp (Figure 6B). As mutations in the *pro35S::Cas9-Psy1* homozygous mutant #1 studied are bi-allelic, result in a premature stop codon and therefore in *PSY1* knock-out (Figure 6C), they lead to uniformly colored fruit peels (Figure 6A). Mutations were detected in both leaf and fruit tissues of *pro35S::Cas9-Psy1* (Figure 6D). In the *pronsLTP::Cas9-Psy1* #4 and #5 mutants studied, ripe fruits were light-red (#4) or light-red/orange (#5) and the color of both fruit flesh and peel was apparently affected (Figure 6A). Consistently, carotenoid accumulation was significantly reduced in ripe fruit pericarp of *pronsLTP::Cas9-Psy1* #4 and #5 (Figure 6B). All the insertion or deletion mutations observed in *pronsLTP::Cas9-Psy1* #4 and #5 resulted in PSY1 proteins truncated in the N-terminal domain of the protein (Figure 6C), which were therefore unlikely to have enzymatic activity. The analysis of mutations frequencies in leaf and fruit tissues indicates that the WT *PSY1* allele was predominant in the leaf, which has only ∼5% of KO *PSY1* alleles detected (Figure 6D). Given that only trichome nuclei are fluorescent in *pronsLTP::NLS-GFP* leaf (Figure 2), the few KO alleles detected in the leaf probably correspond to mutations occurring in the trichomes and have no incidence on the whole plant.

**Figure 6:**
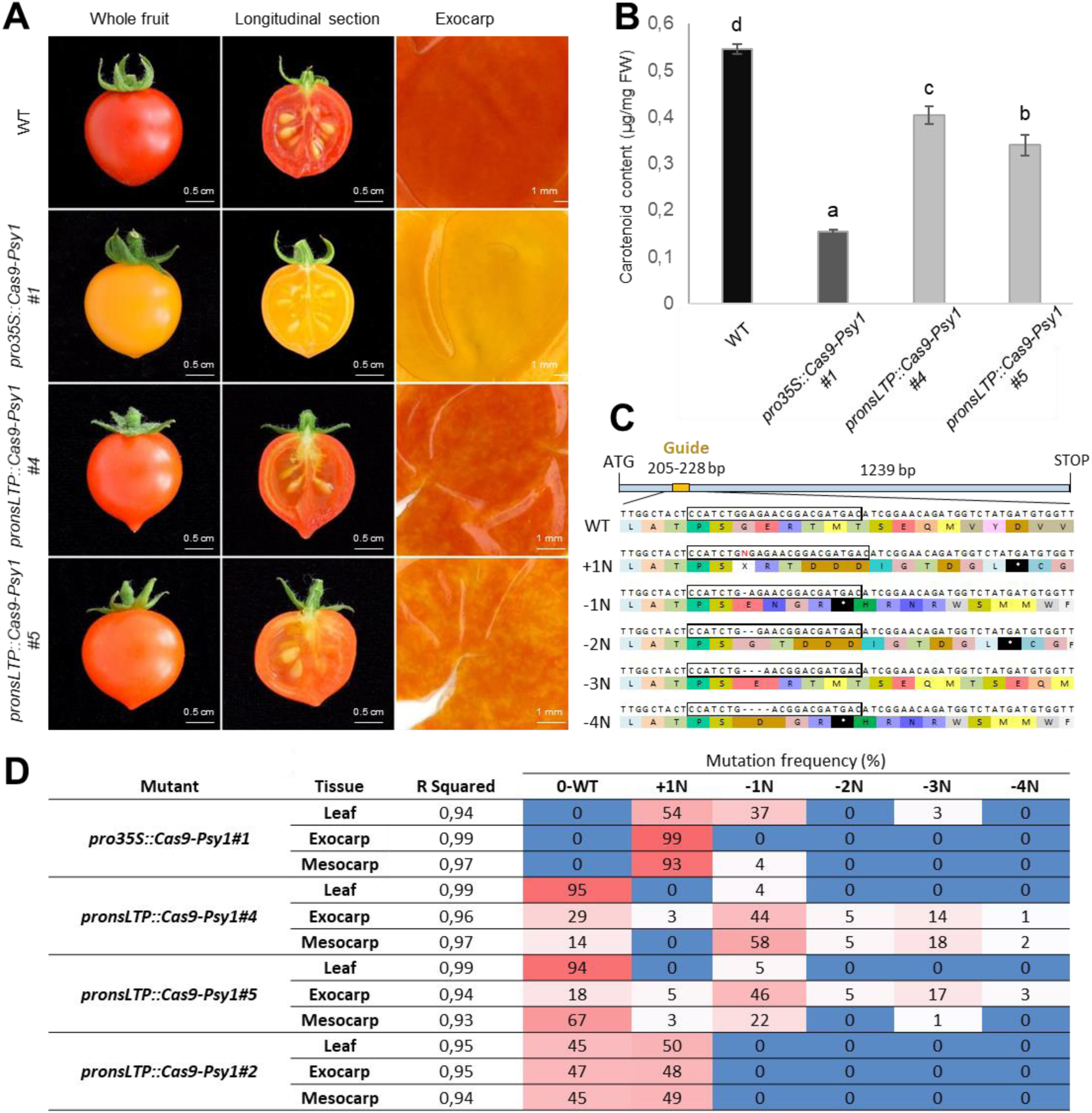
CRISPR/Cas9 mutagenesis of phytoene synthase *SlPSY1* gene using *pronsLTP*. (A) Macroscopic observation of fruit color at RR stage, on whole fruit (left panel), longitudinal section (middle panel) and white light scan of a peeled exocarp (left panel), from WT, *pro35S::Cas9-Psy1* mutant #1, *pronsLTP::Cas9-Psy1* mutant #4 and *pronsLTP::Cas9-Psy1* mutant #5. (B) Carotenoid content of RR fruit pericarp of WT, *pro35S::Cas9-Psy1* mutant #1, *pronsLTP::Cas9-Psy1* mutant #4 and *pronsLTP::Cas9-Psy1* mutant #5. RR (Red Ripe) fruits were collected at Breaker+10. (C) Position of the guide (box) for *SlPSY1* targeting by the CRISPR/Cas9 system. The various mutations detected in *pro35S::Cas9-Psy1* and *pronsLTP::Cas9-Psy1* mutants are indicated below the WT sequence. CRISPR/Cas9-induced frameshift insertion and deletions led to truncated SlPSY1 protein. N, nucleotide; * indicates a stop codon. (D) Frequency and type of mutations observed in leaf and fruit tissues (exocarp, mesocarp plus inner epidermis) of *pro35S::Cas9-Psy1* mutant #1, *pronsLTP::Cas9-Psy1* mutant #4, *pronsLTP::Cas9-Psy1* mutant #5 and *pronsLTP::Cas9-Psy1* mutant #2.

Examination of the fruit peel further revealed its uneven coloration, with light colored yellow to orange or light-red spots appearing on a darker background (Figure 6A). The editing of *PSY1* in exocarp cells likely results in a chimeric peel tissue which collectively displays the range of mutations detected in fruit exocarp of *pronsLTP::Cas9-Psy1* #4 and #5 mutants (Figure 6C, 6D, Figure S5). Individual cell color may range from red (WT alleles or single KO mutations) to light red or yellow-orange due to the accumulation of mono-allelic KO mutations (much reduced PSY1 activity due to haploinsufficiency) or bi-allelic KO mutations (no PSY1 activity). As epidermal cells undergo anticlinal cell divisions^5^, mutations occurring early in fruit development in E1 epidermal cells may be transmitted to daughter epidermal cells, giving to the fruit peel its variegated appearance.

We also detected mutations in the fruit mesocarp tissue of both *pronsLTP::Cas9-Psy1* #4 and #5 mutants (Figure 6C, 6D), whereas the *nsLTP* promoter drives a preferential expression in the fruit exocarp (Figure 2C). A likely explanation is that mesocarp cells derive from exocarp cells, as the two layers of sub-epidermal cells (E2 and E3) undergo periclinal divisions which generate mesocarp cells^5^. While *pronsLTP*-driven expression is much less intense in E2 cells and even less so in E3 cells (Figure 2C) than in E1 cells, and inexistent in mesocarp cells, the detection of KO mutations in fruit mesocarp of *pronsLTP::Cas9-Psy1* #4 and #5 mutants (Figure 6D) suggests that mutated E2 or E3 cells can transmit their mutations to their lineage in mesocarp cells. The fact that the inner epidermis, included in our mesocarp sample, consists of a single cell layer (Figure 2) probably rules out its contribution to the mutations detected in the mesocarp. The identification of a mutant (*pronsLTP::Cas9-Psy1 #2*) carrying the single +1N mutation in a heterozygous state in all tissues analyzed, including the leaf (Figure 6D), suggests that this mutation is inherited and appeared very early in the plant, possibly as early as the transformation and regeneration process.

Thus, we showed here that the *nsLTP* promoter is effective in driving preferential CRISPR/Cas9 mutagenesis in the fruit. The resulting chimeric exocarp tissue carries a set of KO mutations that can be transmitted to the mesocarp, while the plant is not affected. Using the *nsLTP* promoter, we can therefore envisage targeted mutagenesis in the fruit exocarp of single-copy genes that have essential roles in the plant and whose KO mutation can be lethal. An example of these is the *HYDROXYCINNAMOYL-CoA:SHIKIMATE HYDROXYCINNAMOYL TRANSFERASE* (*HCT*) gene, which plays a critical function in the phenylpropanoid pathway and likely in cuticle formation and architecture^12,57^. However, we also observed a mutant carrying an inheritable KO mutation in both fruit and leaf. Our results therefore suggest that, when the aim of using *pronsLTP* is to avoid whole-plant mutagenesis, an additional step of selecting mutation-free plantlets, e.g. after plant regeneration, may be necessary before studying cuticle mutants. To date, one previous study has examined the suitability of the tomato fruit-specific *proPPC2* promoter^29,31^ for fruit-specific gene editing by analyzing phenotypic alterations in plant and fruit due to mutations in a target gene (*ENHANCER-OF-ZEST 2*), and not by sequence analysis^37^, which makes comparison difficult. Likewise, studies on specific CRISPR/Cas9 mutagenesis of various root cell types in *Arabidopsis* could not examine if mutations were also present in non-targeted root cells because of the need of sufficient plant material^58^. Thus, the future development of new tomato promoters for CRISPR/Cas9 mutagenesis of specific fruit, tissue, cell types and/or developmental stage will warrant investigations carefully examining where and when mutations are found in plant and fruit.

## CONCLUSION

Tomato is currently the model for cuticle studies, due to its thick, astomatous, easy-to-isolate cuticle. Tomato cuticle has been instrumental in advancing our understanding of cuticle formation, and particularly of the contribution of cutin embedded polysaccharides, flavonoids and phenolic acids. The question now is to decipher how these compounds modulate cuticle architecture and properties^21,22^. The approach of choice is to modulate their content directly in the cuticle by gene overexpression, silencing (via RNAi) or editing (via CRISPR/Cas9). This strategy aims to avoid any damaging effects on the plant, which can even be lethal when targeting a gene such as HCT, a single phenylpropanoid gene essential to the plant and which probably plays an important role in the cuticle^57^.

To this end, we selected the promoter of the *non-specific Lipid Transport Protein 2* (*pronsLTP*) and showed that, in the fruit, its activity is restricted to the fruit exocarp and inner epidermis, with a strong activity in the outer epidermis at early stages of fruit development, when most of the cuticle material is deposited^24^. We further showed that the *pronsLTP*-driven overexpression of *SlMYB75* induces anthocyanin accumulation specifically in the exocarp of the growing fruit. Using Raman microspectroscopy, we could highlight the resulting alterations in flavonoids and p-coumarate, the main cutin-associated phenolic acid, in the cuticle of both *pro35S-* and *pronsLTP*-driven transgenic lines. However, only *pronsLTP::MYB75* fruits showed significant alteration in fruit properties, including cracking, while changes in cuticle composition were more severe in *prons35S::MYB75* fruits. Altogether, our findings indicate that *nsLTP* can be proposed to the scientific community as a validated promoter suitable for fruit cuticle engineering in tomato. This work also raises apparent discrepancies between the chemical composition of the cuticle and its properties, providing a framework for exploring these relationships. To this end, in-depth investigations will be carried out in the near-future using up-to-date technologies developed for correlative multimodal imaging approach^22^ and the plant material generated here.

We next explored the possibility of using *pronsLTP* for gene editing by CRISPR/Cas9 in the fruit exocarp. To this end, we edited the carotenoid *PSY1* gene with a *pronsLTP*–driven CRISPR/Cas9 system. We detected different knock-out alleles of *PSY1* which were predominantly restricted to the fruit exocarp and mesocarp of color-impaired *pronsLTP::Cas9-Psy1* mutants. Thus, mesocarp cells are also affected, unlike in lines overexpressing *SlMYB75*, which is easily explained by their origin, mesocarp cells deriving from the two sub-epidermal cell layers of the exocarp^5^. Our findings are original in that we were able to show that most of the mutations are fruit-specific, making *pronsLTP* an attractive promoter for gene editing in tomato fruit. They also point out that the *pronsLTP* promoter can be used to efficiently edit cuticle-related genes in the fruit epidermis and, more broadly, provide an example of the feasibility of such approach in fleshy fruits. Given that we also detected a mutant carrying a heritable mutation, and that information on cell/tissue type-specific CRISPR/Cas9 editing in plants remain scarce, the use of the *pronsLTP* promoter, and similar promoters, for gene editing in fruit and epidermis warrants further investigation, for example by targeting other fruit-or epidermis-associated genes.

## Materials and methods

### Plant materials and experimental design

#### Tomato culture conditions

Tomato (*Solanum lycopersicum* cv. Micro-Tom) plants were grown in a greenhouse as described in^59^. Flowers were regularly vibrated to ensure even pollination and tagged at anthesis.

#### Overexpression of *NLS-GFP* and *SlMYB75* and plant transformation

Overexpression vectors were constructed by GoldenGate® cloning of amplified coding sequence of *SlMYB75* (*Solyc10g080610*) and Green Fluorescent Protein in translational fusion to a nuclear localization signal (*NLS-GFP* from the pXK7S*NF2 vector^60^) (see Table S3 for gene specific primers). Open reading frames of *Sl*MYB75 or NLS-GFP were placed under the control of either the *CAMV 35S* promoter (2X35S duplicated promoter) or a 2498 bp DNA fragment located directly upstream of the ATG codon of the *nsLTP2* gene (*pronsLTP*) and followed by OCS terminator and HSP18.2 respectively. A *BpiI* enzymatic restriction site was removed from the *pronsLTP* sequence by 1-bp substitution (C to G) to facilitate cloning. To generate transgenic plants, constructs were transformed into *Agrobacterium tumefaciens* (strain C58C1), and then into tomato (Micro-Tom cv) as previously described^29^ plantlets were further checked for ploidy level by flow-cytometry analysis and polyploid plants were discarded.

#### CRISPR/Cas9-engineered *psy1*

CRISPR/Cas9 mutagenesis of *SlPSY1* (*Solyc03g031860*) was done as previously described^40^ using a single sgRNA guide to induce point mutations in the *PSY1* sequence designed with CRISPOR (http://crispor.gi.ucsc.edu/crispor.py) (Table S3). Resulting mutations were detected in leaves and fruit tissues (exocarp or mesocarp plus inner epidermis) from T1 lines by PCR and Sanger sequencing. Mutations frequency was analyzed with ICE Analysis, 2019, v3.0, from Synthego.

#### Measurement of fruit skin properties

Fruit skin properties were measured as described in^26^. For measurements of cuticle permeability to stain, MG fruits collected from WT, *pro35S::MYB75* and *pronsLTP::MYB75* T2 plants (3 fruits per plant) were placed in 0.1% toluidine blue solution for 16h. After water rinsing, fruit surface staining was visually assessed and photographs were taken under standardized conditions as done in^24^. For water loss measurements, red ripe (RR) fruits harvested from WT, *pro35S::MYB75* and *pronsLTP::MYB75* T2 plants (3 fruits per plant) were stored at 30 °C in a dry oven for 770 hours. Water loss was calculated as a percentage of weight loss. Fruit firmness was assessed by penetrometry using the Fruit Texture Analyser (GüSS, South Africa) equipped with an 8-mm diameter probe, as previously described^40^.

#### Measurement of carotenoid content

Carotenoid concentration was measured from 400 mg of frozen fruit pericarp tissue from T1 plants, ground for 30 seconds using a Dangoumo milling system (Prolabo) at nominal frequency. After acetone extraction^61^, the absorbance of a 50 µL test sample was measured at 474 nm against an acetone blank with an Amersham Ultrospec 3100 Pro (Amersham Bioscience) using a quartz cuvette. The carotenoid concentration, expressed in µg/mg Fresh Weight (FW), was calculated as described in^61^.

#### Macro and Microscopic Analysis

Macroscopic observations of tissues from WT and *pronsLTP::NLS-GFP* T0 plants were carried out on an Axiozoom V16 (Zeiss) using the GFP fluorescence acquisition mode at 496 nm. GFP and white light pictures (in grey level mode) were overlaid to have both signals. For confocal microscopy observations, 3 independent 5 DPA fruits were used for each genotype. Fine sections (120 µm) were obtained using a vibrating blade microtome (Microm HM 650V, Thermo Scientific), stained 30 s in 0.05% calcofluor white (Sigma-Aldrich), washed twice for 5 min with PBS 1X, mounted on a glass slide in PBS 1X and imaged using a laser scanning microscope in confocal mode (Zeiss LSM 880 AIRYSCAN). Photographs of anthocyanin accumulation in plant organs and during fruit development of WT, *pronsLTP::MYB75* and *pro35S::MYB75* T2 lines were taken under standardized conditions. Macroscopic observations in white light were performed with an Axiozoom V16 (Zeiss) on longitudinal sections of 10 DPA fruit. Thick sections (120 µm) of fresh fruit were obtained with a Microm HM 650V vibratome (Thermo Scientific) and imaged with an AxioImager microscope (Zeiss). For confocal acquisition, 150 µm thick sections of 5 DPA fruit pericarp were stained with calcofluor white and observed at 405 nm and 594 nm for anthocyanin autofluorescence with a Zeiss LSM 880 AIRYSCAN. All these experiments were realized at the Bordeaux Imaging Center (http://www.bic.u-bordeaux.fr/).

#### RT-qPCR gene expression analysis

Twelve 1 cm diameter discs of epidermal peel were isolated from six 20 DPA fruits collected from three independent WT, *pronsLTP::MYB75* and *pro35S::MYB75* T2 plants and carefully scratched with a scalpel blade (exocarp samples) as previously described^25^. Two disks per fruit were collected in distinct pools and three pools were made in order to obtain three biological replicates. Pericarp samples, consisting of the whole pericarp minus the peel, were prepared in the same way. Samples were ground in liquid nitrogen and stored at -80°C until RNA extraction. RNA was extracted using the RNA purification from Plant and Fungi Nucleospin kit (Macherey-Nagel). Total RNA integrity and concentration were assessed using an Agilent 2100 Bioanalyzer with RNA Nano Chip (Agilent Technologies). Reverse transcription was carried out using 500 ng of total RNA and Invitrogen SuperScript IV reverse transcriptase following the manufacturer instructions. qRT-PCR was performed with *SlMYB75* specific primers (Table S3) and GoTaq qPCR Master Mix (Promega) using the CFX96 Touch Real-Time system (Bio-Rad). Three biological and three technical replicates were performed per point. *EiF4e* was used as reference gene to calculate the relative expression changes according to the ΔΔCT method.

#### Raman spectroscopy

The cuticles of MG fruits collected from WT, *pro35S::MYB75* and *pronsLTP::MYB75* plants were hand isolated with a spatula and dried before RAMAN analysis. When indicated, cuticles were also submitted to enzymatic purification in baths containing 0.5% cellulase (*Aspergillus sp*., EC 3.2.1.4, Sigma-Aldrich) and 0.5% pectinase (*Aspergillus aculateus*., EC 3.2.1.15, Sigma-Aldrich) during seven days and dried at room temperature. Both hand-isolated and digested cuticles were then used for Raman analysis using an inVia confocal Raman microspectrometer (Renishaw) with a 785 nm laser and a 1200 L/mm grating. Prior to use, the instrument was calibrated using a silicon reference. Spectra were acquired from two independent tomato fruits of each plant on area situated at the junction of three epidermal cells visible with the Raman optical on cuticles before and after enzymatic digestion. This area corresponds to the central furrow of the cuticle commonly observed in cuticle cross-section images.

Measurements were conducted within the range of 300 to 1800 cm^-1^. The laser power was set at 100% and the sample was exposed for six seconds with two accumulations, using a 50× objective (Leica, NA = 0.75). Once the acquisition area had been selected, Z-stack mode was employed to scan the cuticle central furrows in depth, resulting in five acquired spectra on the same location with a 5 µm step interval in the z-axis. This enabled the entire cuticle depth to be scanned. For *pro35S::MYB75* and *pronsLTP::MYB75* fruits, 14 locations were targeted per cuticle on three different pieces of cuticle from the same sample. Of these 14 locations, seven were in white areas (without anthocyanins) and seven in purple areas (with anthocyanins). For WT fruit cuticle and *pro35S::MYB75* and *pronsLTP::MYB75* fruit cuticle after digestion, there were no purple or white areas, so seven different random locations were acquired on three pieces of cuticle. The Raman spectra were pre-processed using the Spectragryph software (optical spectroscopy software, version 1.2.16.1, 2022) with the following steps: smoothing, baseline correction, and area normalization. The average spectra for each sample and acquisition area were plotted for peak attribution. The individual spectra were used for band integration and statistical analysis, with the ANOVA test performed in the Rstudio software.

## Conflict of interest statement

The authors declare that there are no conflicts of interest.

## Author’s contributions

MA, JP, BB and CR conceived and designed the experiments. MA, SP, NG, ADO, JP and EP conducted hands-on experiments and data collection. MA, EP, JP, ML, ADO, DM, BB and CR analyzed the data. CR and BB wrote the original draft. All authors read and approved the final manuscript.

## Acknowledgements

Raman was performed at the PROBE research infrastructure, Biopolymers Interactions, Structural Biology (BIBS) facility at Nantes. We thank JP Mauxion for the initial cloning of the *nsLTP* promoter and N Bollier for helpful discussions on the CRISPR/Cas9 system. This work was supported by the ANR (Agence Nationale de la Recherche) grant COPLAnAR (ANR-21-CE11-0035). MA fellowship was supported by COPLAnAR. EP was supported by a PhD fellowship granted by COPLAnAR and a grant “Soutien Trajectoire” from Region Pays de la Loire.

## Data and material availability

All relevant data generated or analyzed are included in the manuscript and the supporting materials. The materials (e.g. vectors) that support the findings of this study are available on request from the corresponding author.

## Supplementary data

**Figure S1.** Wax, cutin, phenolic and flavonoid biosynthetic pathways in the fruit epidermis.

**Figure S2.** The *nsLTP2* co-expressed gene cluster involved in flavonoid biosynthesis.

**Figure S3.** Types of trichomes found on the leaf surface of *S. lycopersicum* var. Micro-Tom.

**Figure S4.** Raman analysis of enzymatically isolated cuticle from tomato fruit.

**Table S1.** Digital expression of tomato genes in the outer epidermis of the fruit.

**Table S2.** Digital expression of tomato genes in fruit tissues along development.

**Table S3.** List of primers used in the study.

## Notes

### Competing Interest Statement

The authors have declared no competing interest.

