## Supplementary material for "Targeting the tomato fruit cuticle by gene overexpression and editing with the fruit epidermis-preferential *nsLTP* promoter": Sup Table S3

**Supplemental Table S3 List of primers used in the manuscript**.

| Name for paper | Forward 5’-3’ | Reverse 5’-3’ | Use |
| --- | --- | --- | --- |
| pronsLTP | GACCTTTCTGAAAGATTGTGA | TCTTATAAAAAAATTGAGTAAAGATTATAGAT | Cloning |
| pronsLTP-mut-BpiI | TGTCTTTTGTCTTTCTCTTTCTCGC | GACACCTAAGCATGAGAATATTTGGA | Cloning |
| *SlMYB75* | TTGAAGACAAAATGAATACTCCTATG | TTGAAGACAAAAGCTTAATTAAGTAG | Cloning |
| SlMYB75-qPCR | ACTTCCAGGAAGGACAGCAAA | GACGAGGATGAGAACGAGGA | qRT-PCR |
| NLS-GFP | TTGAAGACAAAATGCAGCCTTCTCTTAAACGCAT | TTGAAGACAAAAGCTTACTTGTACAGCTCGTCCATGC | Cloning |
| SlPSY_guide |  | TCATCGTCCGTTCTCCAGA | Cloning |
